# A powerful framework for an integrative study with heterogeneous omics data: from univariate statistics to multi-block analysis

**DOI:** 10.1101/357921

**Authors:** Harold Duruflé, Merwann Selmani, Philippe Ranocha, Elisabeth Jamet, Christophe Dunand, Sébastien Déjean

## Abstract

The high-throughput data generated by new biotechnologies used in biological studies require specific and adapted statistical treatments. In this work, we propose a novel and powerful framework to manage and analyse multi-omics heterogeneous data to carry out an integrative analysis. We illustrate it using the package mixOmics for the R software as it specifically addresses data integration issues. Our work also aims at confronting the most recent functionalities of mixOmics to real data sets because, even if multi-block integrative methodologies exist, they still have to be used to enlarge our know-how and to provide an operational framework to biologists. Natural populations of the model plant *Arabidopsis thaliana* are employed in this work but the framework proposed is not limited to this plant and can be deployed whatever the organisms of interest and the biological question. Four omics data sets (phenomics, metabolomics, cell wall proteomics and transcriptomics) have been collected, analysed and integrated in order to study the cell wall plasticity of plants exposed to sub-optimal temperature growth conditions. The methodologies presented start from basic univariate statistics and lead to multi-block integration analysis, and we highlight the fact that each method is associated to one biological issue. Using this powerful framework led us to novel biological conclusions that could not have been reached using standard statistical approaches.

## 1. INTRODUCTION

Biological processes can be studied using measurements that are ever more complex. Today, biologists have access to plethora of new technologies to address their questions. The high-throughput measurements have revolutionized the way to evaluate and predict the behavior of organisms for example in response to environmental changes. Nowadays, one biological sample can deliver many types of “big” data, such as genome sequences (genomics), genes and proteins expression levels (transcriptomics and proteomics), metabolite profiles (metabolomics) and phenotypic observations (phenomics). The revolution of high throughput technologies has also greatly reduced the cost of those omics data production, opening new prospects to the development of tools for data treatment and analysis (Li, Wu, & Ngom, 2016; Meng et al., 2016).

The heterogeneous data collected from cellular to organism levels are associated to a wide variety of techniques sometimes species-specific. The acquisition of data requires a particular experimental design and a suitable methodology to highlight their mining (Rai, Saito, & Yamazaki, 2017). An experimental design inadequate for an integrative analysis could complicate the final interpretation of the collected data. On the contrary, a suitable methodology of analysis can be optimized and brings keys to improve the visibility of the whole data. This point of view was previously stated in (Kerr, 2003) for microarray studies: “*While a good design does not guarantee a successful experiment, a suitably bad design guarantees a failed experiment—no results or incorrect results*”.

Use of multi-omics data makes possible a deeper understanding of a biological system (Zargar et al., 2016, Rajasundaram & Selbig, 2016). Indeed, quantification technologies improve accuracy and create great potential for elucidating new questions in biology. However, this technological revolution must be carefully used because the correlation between quantification analyses is not effective. For example, it is known that it is usually difficult to correlate transcriptomic and proteomic data (Duruflé et al., 2017; Jamet et al., 2009; Maier, Güell, & Serrano, 2009). Each of these technologies has its own limitations and collecting different types of data should help understanding the effects of one or more experimental conditions. A cohort of hypotheses can be proposed with multi-omics analysis. Thus, biological candidates can be identified as biomarkers (*e.g.* genes, proteins, molecules) under complex environmental conditions, and/or new complex regulations can be found.

Altogether, it is generally admitted that studying a single kind of omics data is not sufficient to understand the effects of a treatment on a complex biological system. To obtain a holistic view, it is preferable to combine multiple omics analyses. To highlight the interest of such integrative approaches, let us consider a toy example with two variables (Vx and Vy) measured on 12 individuals (6 from one group called Controlled, and 6 from another one called Treated). The values are presented in supplemental Table S1. Statistical tests (Wilcoxon rank sum test and Student t test) do not reveal any significant difference between the two groups for both the Vx and Vy variables when they are analysed separately (p-values higher than 0.3). But, a simple scatterplot (Figure 1) highlights the interest of combining the two variables. Indeed, it clearly appears that the two groups are separated if we consider the Vx and Vy variables together.

**Figure 1.**
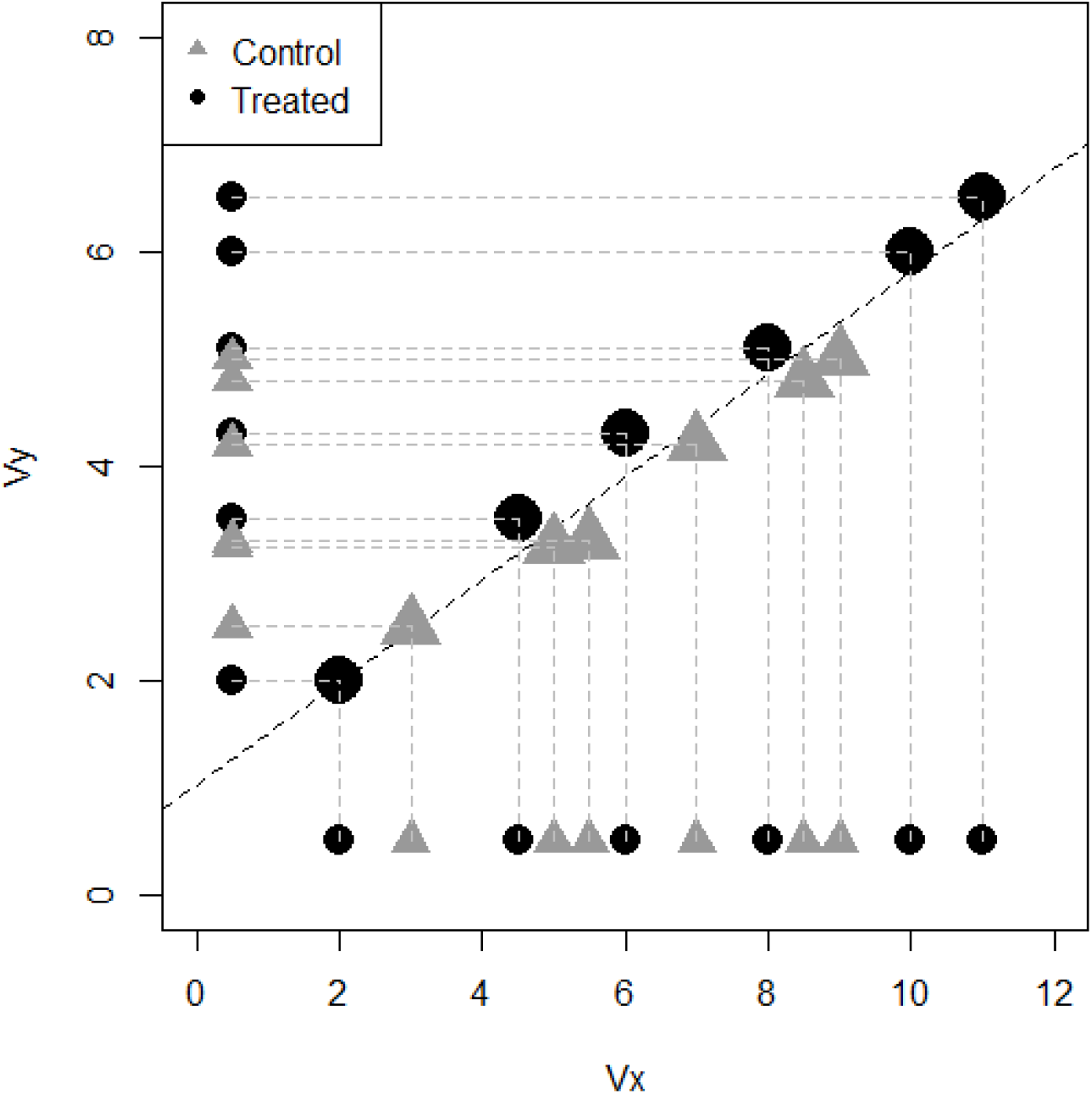
Scatterplot representing Vy values (vertical axis) according to Vx values (horizontal axis). Control and treated observations are represented with grey triangles and black circles, respectively.

Thus, in the same vein, we claim that the integrative analysis of several data sets acquired on the same individuals can reveal information that single data set analysis would keep hidden. Furthermore, the toy example also highlights the interest of a relevant graphical representation: information hidden in supplemental Table S1 is clearly visible in Figure 1. The recent work by (Matejka & Fitzmaurice, 2017) is assuredly a good way to be strongly convinced about data visualisation.

This article focuses on a powerful framework we propose to manage and analyse heterogeneous data sets acquired on the same samples. It proceeds step by step, from basic univariate statistics to multi-block integration analysis (Singh et al., 2016; Tenenhaus et al., 2014). We illustrate the gaps bridged by each method from the computation of univariate statistics to a thorough implementation of multi-block exploratory analysis. The implementation of the methods and the graphical visualizations have simply been accomplished with existing tutorials for the R software (R Core Team, 2018) and the mixOmics package (Rohart et al., 2017). But, since their interpretation is not easy (González et al., 2013), this article will provide a better understanding of the statistical integration and a way to include it in a global reflexion structured in a workflow summarized in Figure 2. We also aim at increasing our know-how related to these novel methodologies by confronting them to new real data sets. The first section presents the background of our study and the data sets we have dealt with detailed in (Duruflé, 2019a; 2019b). Then, we describe several statistical methods used to address specific biological questions. Afterwards, we explain in detail the statistical results and give clues to interpret them.

**Figure 2.**
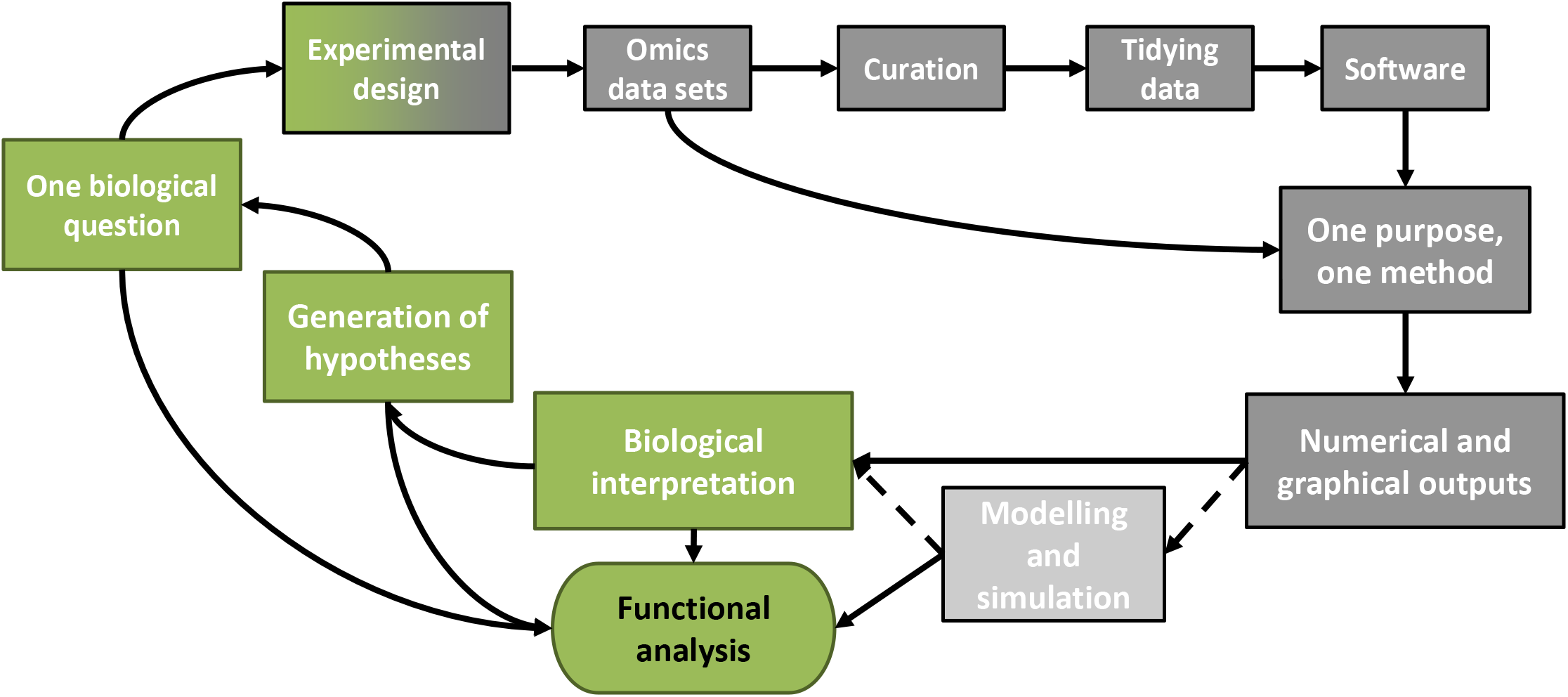
Workflow for our multi-omics integrative studies. The different parts of this article are represented with grey boxes and the green boxes close the workflow with biological concepts. The workflow converges towards the functional analysis required to validate the whole study.

## 2. BIOLOGICAL CONTEXT

In the global warming context, seasons are altered with modifications of the temperatures. The elevation of the temperature is the most studied change because it is already observed (Savo et al., 2016). The occurrence of cold stress can also appear without any previous chilling period and it could become a problem to maintain agricultural productivity in the future (Gray & Brady, 2016). The model plant *Arabidopsis thaliana* of the *Brassicaceae* family has a worldwide geographical distribution and therefore has to adapt to multiple and contrasted environmental conditions (Hoffmann, 2002). The huge accumulation of molecular data concerning this plant is very helpful for studying complex multiple levels responses. It is expected to transfer obtained results to other plant species of economic interest for translational pipelines (Sibout, 2017).

### 2.1. Experimental design

First, a compromise is necessary to determine the ideal number of biological replicates. It is hard to find an agreement between the reality of the biological experimentation (*e.g.* limitation in material, space, time, work force and cost) and the necessity to get robust information for the statistical analyses. The method used for the randomization of the replicate also needs to be considered. For these reasons, the experimental protocol must minimize potential external impacts within and between the replicates and avoid confounded effects.

To strengthen the results, each biological replicate can be the average of several technical replicates, if the type of analysis allows it. For the biologist, it is important to know the number of experimental repetitions to appreciate the variability between the different conditions. But, for a statistician, the information resides into the intrinsic variability of the different samples or repetitions. For all these reasons, one sample considered as “out of norms” by the biologist could be valuable in a multi-omics analysis.

Our experimental design was built with two crossed factors: i) ecotypes with 5 levels (4 Pyrenees Mountain ecotypes Roch, Grip, Hern, Hosp, living at different altitudes, and Col, a reference ecotype from Poland, living at low altitude) and ii) temperature with 2 levels (22°C and 15°C). For each ecotype, rosettes and floral stems were collected and analysed. At 22°C, rosettes were collected at 4 weeks, *i.e.* at the time of floral stem emergence. At 22°C, floral stems were collected at 6, 7 and 8 weeks respectively for Col, Roch / Grip and Hern / Hosp. At 15°C, rosettes and stems were collected 2 weeks later than at 22°C. More details about the plant culture conditions can be found in (Duruflé, 2019a). Three independent biological replicates were analysed for each sample including 20 plants per sample. To minimize the experimental effect, each plant was grown at a randomly chosen place according to the experimental design represented in Figure 3.

**Figure 3.**
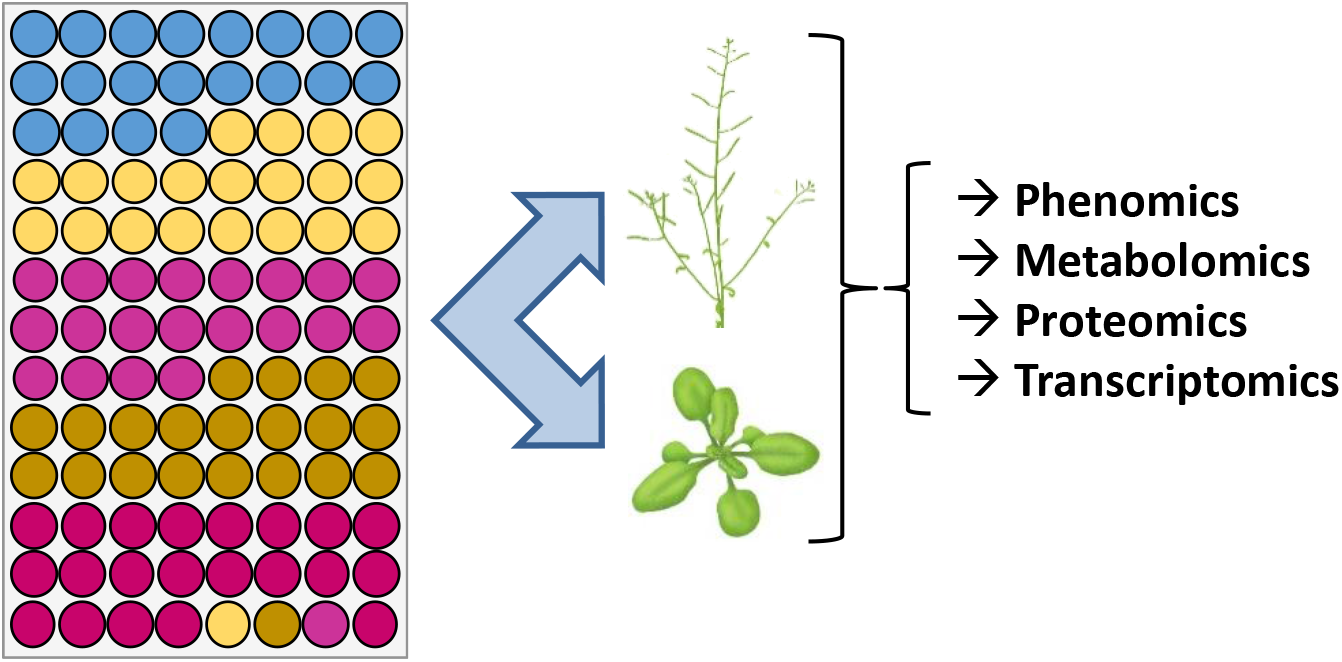
Schematic overview of the strategy and experimental protocol used in this study. Each circle represents one plant and each color stands for one ecotype of *A. thaliana*. For each of the three biological replicates, the position of a given ecotype has been changed randomly to avoid position effects.

### 2.2. Omics data sets and curation

In this project, the four following omics data sets (called blocks thereafter) were collected:

i. Phenomics, *i.e.* a macro phenotyping analysis, was performed on two organs: rosettes and floral stems (Duruflé, 2019a). Indeed at the time of sample collection and prior to freezing, 9 phenotypic variables were measured: 5 on rosettes (mass, diameter, number of leaves, density, and projected rosette area), and 4 on floral stems (mass, diameter, number of cauline leaves, length).
ii. Metabolomics, *i.e.* identification and quantification of seven cell wall monosaccharides (fucose, rhamnose, arabinose, galactose, glucose, xylose and galacturonic acid), were performed as previously described (Duruflé et al., 2017). Theoretical cell wall polysaccharide composition was inferred, based on the monosaccharide analyses according to (Duruflé et al., 2017; Houben et al. 2011; Duruflé, 2019a).
iii. Proteomics, *i.e.* identification and quantification of cell wall proteins by LC-MS/MS analyses, were performed as described (Duruflé et al., 2017). Altogether, 364 and 414 cell wall proteins (CWPs) were identified and quantified in rosettes and floral stems, respectively (Duruflé, 2019b).
iv. Transcriptomics, *i.e.* sequencing of transcripts also called RNA-seq, was performed according to the standard Illumina protocols as described (Duruflé et al., 2017). Altogether, 19763 and 22570 transcripts were analysed in rosettes and floral stems, respectively (Duruflé, 2019b).

## 3. TIDYING DATA

Statistical data analysis requires efficient data pre-processing. As mentioned in (Wickham, 2014), “*It is often said that 80% of data analysis is spent on the process of cleaning and preparing the data”*. So in an integrative analysis framework, each data set needs to be structured in the same way, and (Wickham, 2014) has also stressed the following statements: *1/ Each variable forms a column. 2/ Each observation forms a row*. So, in our context, each data set is structured with biological samples in rows and variables in columns.

Handling missing data is always a big deal. As stated by Gertrude Mary Cox (an American statistician of the 20^th^ century), “*the best thing to do with missing values is not to have any”*. Fortunately, many methods exist to deal with missing values. For instance, the methodologies implemented in the missMDA package (Husson & Josse, 2013) are dedicated to the handling of missing values in the context of multivariate data analysis. For example in this work, missing proteomics quantification data were dealt with considering two situations: (i) non-validated proteins (identification with a single specific peptide and/or in a single biological replicate); and (ii) undetectable proteins (no peptide identified in a given condition). In the former case, a background noise, corresponding to the minimum, and the first statistical quartile of the biological replicate, was applied. In the latter case, a background noise of 6 (value lower than the minimum value found in the whole experiment) was applied. This treatment allowed combining the quantification process with the qualitative study and provided a higher confidence in the final result.

More recently, a study focused on missing rows in data sets in an integrative framework (Voillet et al., 2016). Within an integrative study, we can easily be in this case if, for instance, the number of biological replicates is not the same for transcriptomics and proteomics analyses. The main idea to remember would be to deal with missing values with an *ad-hoc* method taking into account the specificity of the data. In our case, two replicates of the transcriptomic data had to be deleted due to their low quality. Following the method proposed in (Voillet et al., 2016), these missing rows were imputed using the samples for which all the data were available, *i.e.* the two other replicates.

## 4. RATIONALE SUPPORTING THE PROPOSED FRAMEWORK

### 4.1. Software

As mentioned in the Comprehensive R Archive Network (CRAN, cran.r-project.org), *R “is a freely available language and environment for statistical computing and graphics which provides a wide variety of statistical and graphical techniques: linear and nonlinear modelling, statistical tests, time series analysis, classification, clustering, etc*.”

R functions with a command-line interface that, even if it can appear not user-friendly, allows the user to build scripts that can be run on various data sets with rather few tuning. R gives access to the newest methodological developments due to its very active community (R-bloggers, R-help, UseR conference…) motivated by open science considerations. Furthermore, efficient tools such as RStudio (www.rstudio.org) were developed in order to make the initiation to R easier. In addition, many resources are available on CRAN to start with R. Therefore, it seems highly reasonable to expect that the user can read, use and adapt existing scripts available in the examples of each manual of packages after few hours of practice. Specifically considering the community of biologists using R, the Bioconductor repository (http://www.rstudio.org/) provides selected tools for the analysis of high-throughput genomic data (Gentleman et al., 2004).

The dynamism around R appears in the packages developed by and for the community. So, several packages exist to address statistical integrative studies. We focus on the mixOmics package (Lê Cao et al., 2009; Rohart et al., 2017), but other packages such as FactoMineR (Husson & Josse, 2013) can also be used for nearly similar purposes. Methodologies presented in (Bécue-Bertaut & Pagès, 2008) and (Sabatier et al., 2013) are also alternatives, as well as the Multi-Omics Factor Analysis (MOFA) approach proposed in (Argelaguet et al., 2018). Regarding commercial software for instance, SIMCA-P (Umetrics, umetrics.com/) propose several methods to perform integrative analyses, and toolboxes for Matlab are also available (The MathWorks, Inc., Natick, Massachusetts, United States). We choose to favor an open source software, as it is easier to promote a free software than a commercial one when people are not specialists in the domain (Carey & Papin, 2018). Furthermore, mixOmics appears as a very active package addressing data integration issues. It has been downloaded more than 25,000 times (unique IP address) in 2017, 5 versions were released in 2017, the reference article (Lê Cao et al., 2009) has been cited 300 times and the mixOmics team has published 16 articles related to this package since 2008.

### 4.2. One purpose, one method

In this section, partly inspired from the tutorial of mixOmics (mixomics.org), we wish to highlight the link between a biological question (purpose) and the appropriate statistical method.

- *Purpose: explore one single quantitative variable (e.g. what is the level of expression of one gene?).* Method: univariate elementary statistics such as mean, median for main trends, and standard deviation or variance for dispersion, can be completed with a graphical representation such as boxplot.
- *Purpose: assess the influence of one single categorical variable on a quantitative variable (e.g. Are the plant growth different in two or more environmental conditions?)*. Method: statistical significance test such as Student t test or Wilcoxon rank sum test for two groups and ANOVA or Kruskall-Wallis for more groups will address this question (McDonald, 2009). In this context, a special attention must be paid to the structure of the data: independent samples (*e.g.* independent groups observed in various conditions) or paired samples (*e.g.* same samples observed twice or more in various conditions).
- *Purpose: evaluate the relationships between two quantitative variables (e.g. Is there a correlation between the concentration of one protein and its transcript abundance?).* Method: correlation coefficients (Pearson for linear relationships and Spearman for monotonous ones) (McDonald, 2009). Graphical representations of correlation matrices can provide a global overview of pairwise indicators (Friendly, 2002; Murdoch & Chow, 1996).
- *Purpose: explore a single data set (*e.g. *transcriptomics) and identify the trends or patterns in the data, experimental bias or, identify if the samples ‘naturally’ cluster according to the biological conditions (e.g. Can we observe the effect of different environmental growth conditions on different ecotypes?)*. Method: an unsupervised factorial analysis such as Principal Component Analysis (PCA) (Mardia, Kent, & Bibby, 1980) provides such information about one data set without any *a priori* on the result. Centering and scaling the data, such that all variables have zero mean and unit variance, before performing PCA is usually useful when dealing with omics data to make the PCA results meaningful.

The previously mentioned methods are rather standard and usually used for biological data analysis whereas the methods mentioned hereafter are less usual.

- *Purpose: classifying samples into known classes based on a single data set (e.g. Can we classify various ecotypes according to their transcriptomics profile?)*. Method: supervised classification methods such as Partial Least Square Discriminant Analysis (PLS-DA) (Lê Cao, Boitard, & Besse, 2011) assess how informative the data are to rightly classify samples, as well as to predict the class of new samples.
- *Purpose: unravel the information contained in two data sets, where two types of variables are measured on the same samples (e.g. What are the main relationships between the proteomics and transcriptomics datasets?).* Method: using PLS-related methods (Wold et al., 2001) enable knowing if common information can be extracted from the two data sets (or highlight the relations between the two data sets).

The following methods are very recent and few applications have been published so far. This work contributes to improve their efficiency on real data sets.

- *Purpose: the same as above but considering more than two data sets (e.g. What are the main relationships between the proteomics, transcriptomics and phenotypic data?)*. Method: multi-block PLS related methods were recently developed to address this issue (Günther et al., 2014; Singh et al., 2019).
- *Purpose: the same as above but in a supervised context (e.g. Can we determine a multi-omics signature to classify ecotypes?)*. Method: multi-block PLS-DA (referred as DIABLO for Data Integration Analysis for Biomarker discovery using Latent variable approaches for Omics studies) was recently developed to address this issue (Singh et al., 2019).

A schematic view of the data sets and the methods implemented is presented in Figure 4. The way to perform an integrative statistical study is illustrated through several cycles (Figure 4B). We prefer this view rather than a straightforward pipeline beginning with univariate analysis and ending with multi-block approaches. Each method contributes to the global comprehension of the data and can challenge the others. For instance, univariate statistics may highlight outliers or essential variables. On the other hand, multi-block approaches may focus on new samples and/or variables showing specific behavior that should be studied through a univariate method. We claim that, facing integrative studies, a relevant statistical analysis must go through these cycles, with progress and feedback.

**Figure 4.**
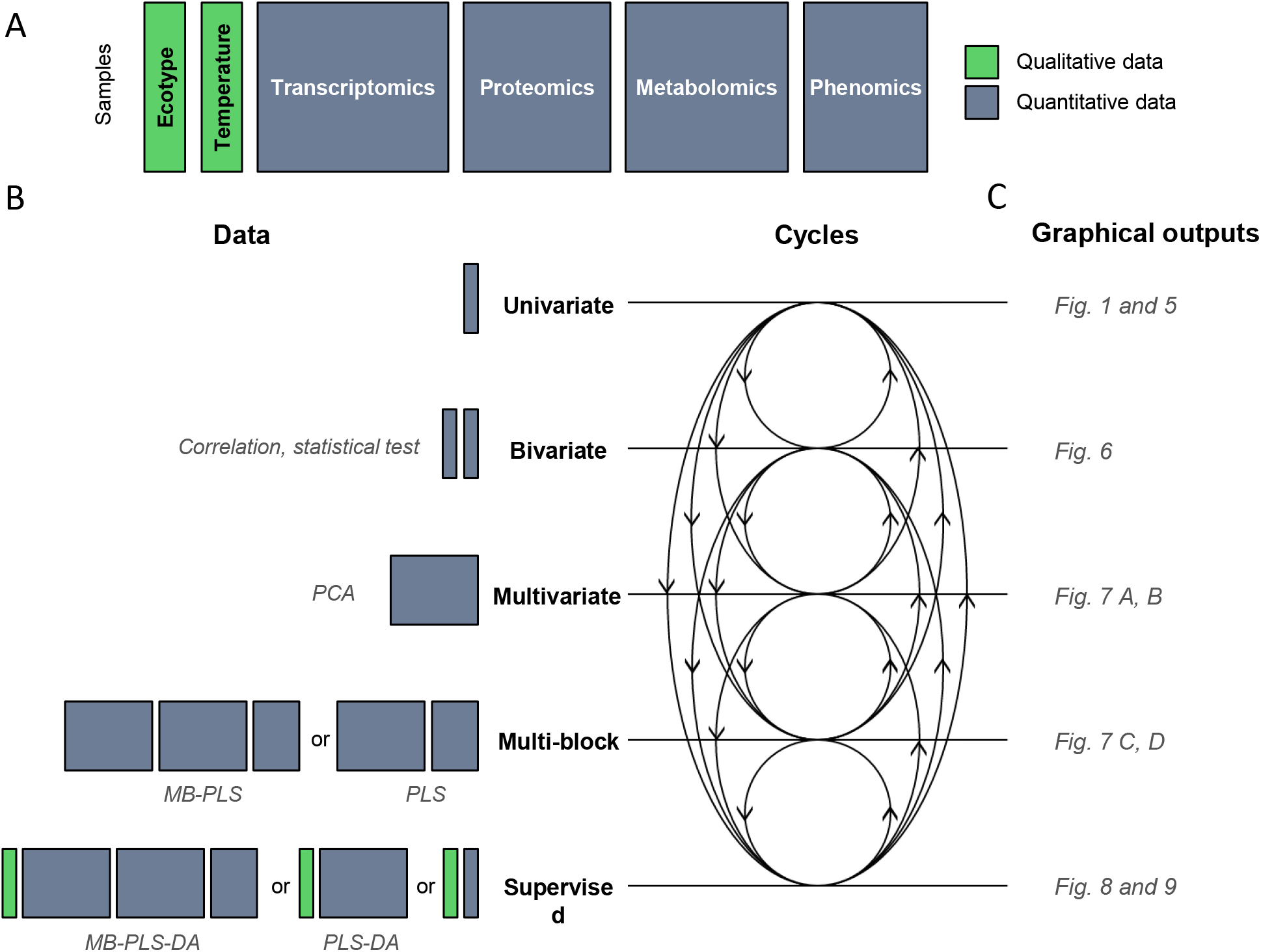
One purpose, one method to analyse qualitative and quantitative blocks. A) Schematic representation of the different blocks (or data sets) co-analysed in this study. The samples are represented in rows and the variables in columns. B) Schematic overview of the methods implemented represented by cycles within an integrative study. C) Examples of graphical outputs detailed in the results section. PCA: Principal Component Analysis; MB: Multi-Blocs; PLS: Partial Least Squares regression; DA: Discriminant Analysis. Qualitative and quantitative blocks are represented in green and grey respectively.

### 4.3. Sparse extensions

Every methods developed in mixOmics are proposed with a sparse extension (sparse PCA (S-PCA), sparse PLS (S-PLS)…). Sparse methods are useful to remove non-informative variables (*e.g.* which can be considered as background noise) regarding the purpose of the multivariate method. Concerning PCA for instance, the sparse version selects only the variables that highly contribute to the definition of each principal component (PC), removing the others. Sparsity is mathematically achieved via Least Absolute Shrinkage and Selection Operator (LASSO) penalizations (Tibshirani, 1996).

In practice, the use of sparse methods in the context of omics data is very useful as it reduces the number of potentially relevant variables displayed on the graphical outputs. Thus, it facilitates the biological interpretation of the results and minimizes the list of potential candidates for further investigations.

### 4.4. Numerical and graphical outputs

As previously mentioned, statistical analysis should be associated with graphical representations (Figure 4C). A famous sentence assigned to Francis John Anscombe (a British statistician of the 20^th^ century) emphasized this point of view: “… *make both calculations and graphs. Both sorts of output should be studied; each will contribute to understanding.” (Anscombe, 1973)*. Based on this principle, a recent work by Matejka and Fitzmaurice (Matejka & Fitzmaurice, 2017) illustrates in a quite funny way how same numerical outputs can provide very different graphical representations (including a scatterplot looking like a dinosaur named *datasaurus*).

The results of univariate and bivariate approaches are mainly reported as p-values for statistical testing. Boxplots and barplots, as produced, for instance, by the ggplot2 package (Wickham, 2016), may complete and reinforce the interpretation of the results (Figure 4C). Regarding barplots, one core question relies on the error bars that are frequently added: should they be based on standard deviation or on standard error of the mean? A thorough explanation about the difference is provided in (Cumming, Fidler, & Vaux, 2007). The authors also mention this statement that may seem obvious but that is sometimes forgotten: “*However, if n is very small (for example n = 3), rather than showing error bars and statistics, it is better to simply plot the individual data points.”*

We also used graphical representations of correlation matrices (Figure 4C) such as those produced by the corrplot package (Wei & Simko, 2016) for the R software. This is essential when dealing with (not so) many variables: with 50 variables, 1225 (50 × 49 / 2) pairwise correlation coefficients are computed and have to be analysed and interpreted.

Regarding multivariate analyses (from PCA to multi-block analyses), we used the graphical outputs provided by the mixOmics R package (Rohart et al., 2017). They are based on the representation of individuals and variables projected on specific sub-spaces (Figure 4C). A thorough discussion about the complementarity between several graphical displays is given in (González et al., 2013).

In a multivariate supervised analysis, the individuals (biological samples) of the study are represented as points located in a specific sub-space defined by the first PLS-components (Figure 4C). Interpretation is based on the relative proximities of the samples and on the equivalent representation for variables.

The standard representation for the variable plots is frequently referred as correlation circle plot (Figure 4C). It was primarily used for PCA to visualise relationships between variables, but it has been extended to deal with multi-block analysis. In such a plot, the correlation between two variables can be visualised through the cosine of the angle between two vectors starting at the origin and ending at the location of the point representing the variable. The representation of variables can also be done through a relevance network. These networks are inferred using a pairwise similarity matrix directly obtained from the outputs of the integrative approaches (González et al., 2013). A Circos plot (Singh et al., 2019) can be viewed as a generalization of relevance network where the nodes are located on a circle. Then, based on the same pairwise similarity matrix used for relevance network, a clustered image map can be displayed. This type of representation is based on a hierarchical clustering simultaneously operating on the rows and columns of a real-valued similarity matrix. This is graphically represented as a 2-dimensional colored image, where each entry of the matrix is colored on the basis of its value, and where the rows and columns are reordered according to the hierarchical clustering.

## 5. RESULTS

In this section, we provide neither a thorough biological interpretation of the results, nor a comprehensive view of every statistical analysis performed. Instead, we highlight the limits of each method leading to the next step of the statistical analysis and show how a biologist can interpret and take over the conclusions of a statistical study.

### 5.1. Bivariate analysis

We illustrate the bivariate analysis through some graphical representations of phenotypic data linked to one parameter of the experimental design. Figure 5A displays parallel boxplots as well as individual observations of the number of leaves for the 5 ecotypes at the 2 growth temperature conditions. Figure 5B only displays the average values of one triplicate for each ecotype and temperature.

**Figure 5.**
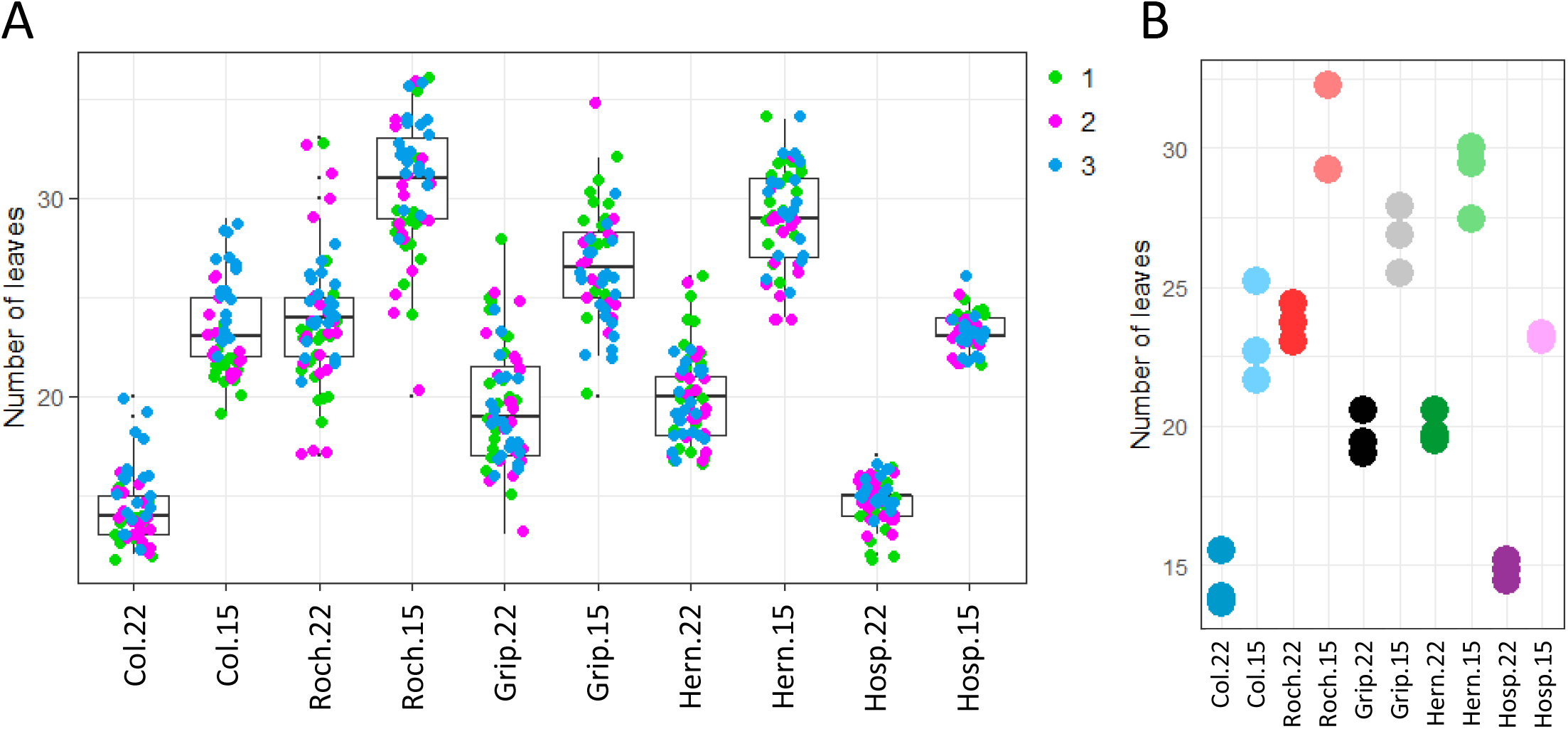
Examples of graphical outputs of a supervised bivariate analysis illustrated by A) A boxplot (each color corresponds to the different values obtained for each triplicate) and B) An individual plot. (each color corresponds to the average obtained for one triplicate, and does not match with color used in A). The number of leaves for 5 ecotypes of *A. thaliana* (Col, Roch, Grip, Hern and Hosp) and 2 growth temperatures (22 and 15°C) was used. These plots were obtained using functions geom_point() and geom_boxplot() from the ggplot2 package (Wickham, 2016).

The main information extracted from these graphics concerns a quality control of the data. The relatively low scattering of points representing individuals of each biological replicate (Figure 5A) indicates a rather good reproducibility between all the samples and between the repetitions. So, the values from several plants of a given biological repetition can be averaged, to go on with the analyses. The visual impression provided by Figure 5B regarding the temperature and ecotype effects can be confirmed via statistical testing such as two-way ANOVA (Bingham & Fry) (results not shown). However, this kind of analysis does not provide any information about the potential relationships between several variables. This drawback justifies the next step of analysis which deals with a whole data set.

### 5.2. Multivariate analysis

The multivariate approach is illustrated on the rosettes cell wall transcriptomics data set. It is composed of 364 variables (or transcripts). The first way to question the whole data set can be through the computation of pairwise correlation coefficients. For instance, Figure 6 displays the correlation matrix between samples. It indicates that the levels of gene expression for each sample are positively correlated (only green color and identically oriented ellipses) with all the others.

**Figure 6.**
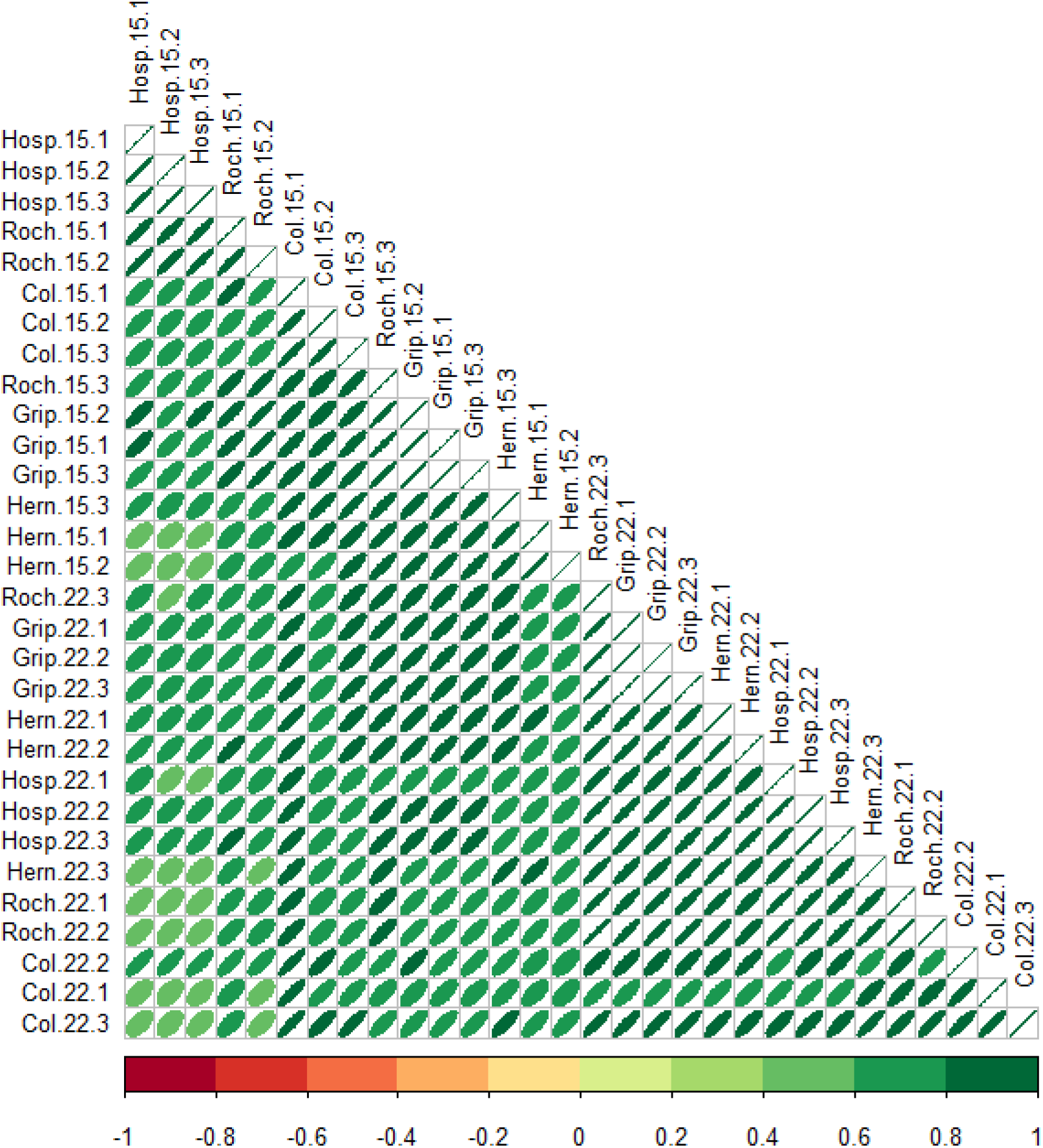
A graphical representation of the multivariate analysis, pairwise correlation coefficients of cell wall transcriptomics data sets in the rosettes of the five *A. thaliana* ecotypes grown at 15°C or 22°C. The color code and the ellipse size represent the correlation coefficient between the levels of expression of genes for each sample. The areas and the orientations of the ellipses represent the absolute value of the corresponding correlation coefficients. The eccentricity of the ellipses represents the absolute value of the corresponding correlation coefficients. This plot was obtained using the function corrplot() from the corrplot package (Wei & Simko, 2016).

Then, a PCA can be performed as an extension of the quality control. For instance, Figure 7A highlights the distance between the three replicates corresponding to one condition. We can observe that the Grip ecotype is well gathered, whereas the Col ecotype is more scattered. This information must be moderated because of the rather low proportion of variance explained by the first two principal components displayed here. Having a look at the following components could be meaningful to consolidate and complete this information.

**Figure 7.**
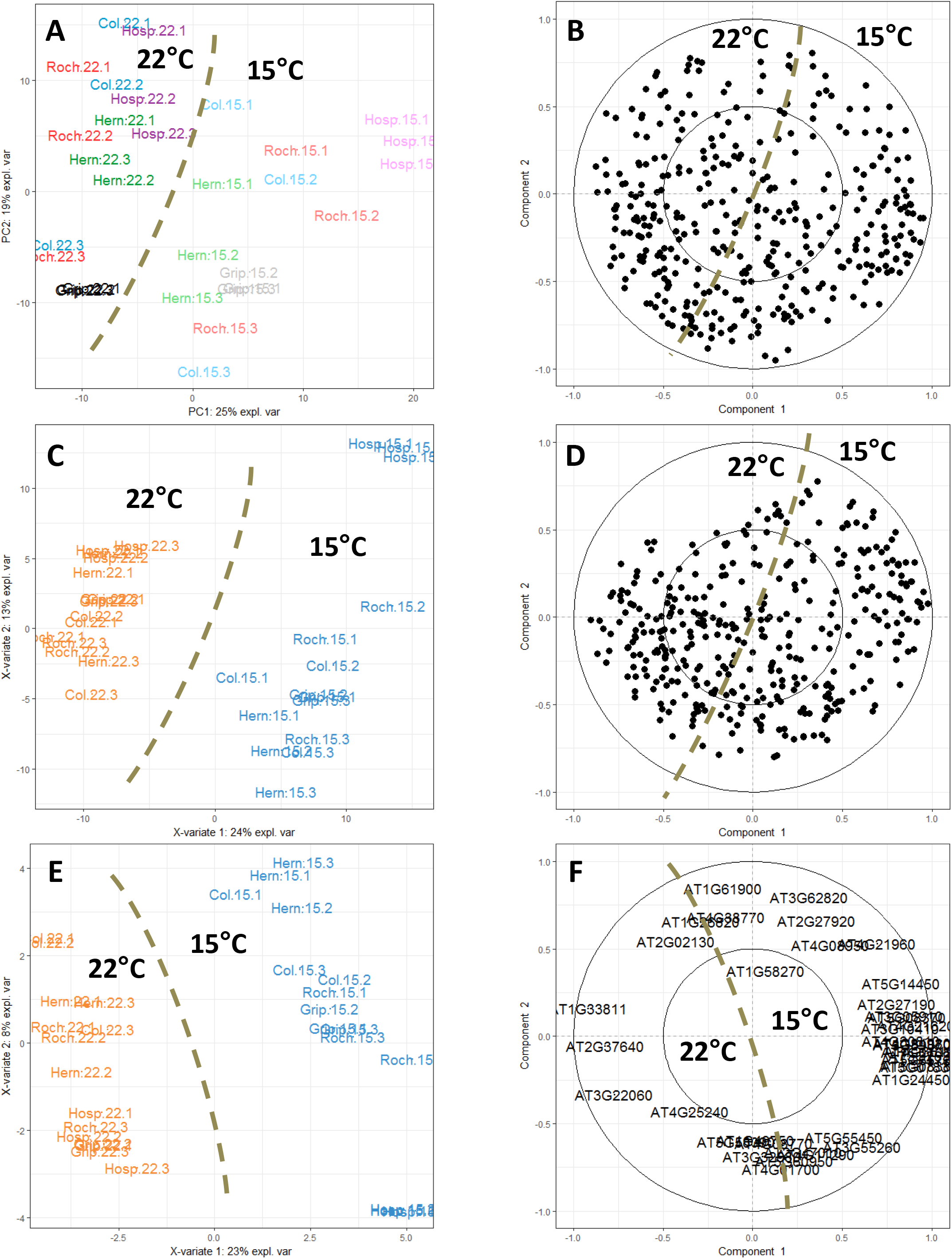
Graphical representation of the unsupervised (A, B) and supervised (C-F) analysis of the rosette cell wall transcriptomes from ecotypes grown at 22°C and 15°C. A) Individuals plot of a PCA from ecotypes grown at 22°C (bright color) and 15°C (pale color) associated to the B) Variables plot. C) Individuals plot of a PLS-DA from ecotypes grown at 22°C (orange) and 15°C (blue) associated to the D) Variables plot and E) Individuals plot of a S-PLS-DA associated to the E) Variables plot. Two circles of radius 1 and 0.5 are plotted in each variables plot to reveal the correlation structure of the variables. These plots were obtained using the functions pca(), plsda(), plotIndiv() and plotVar() from the mixOmics package (Rohart et al., 2017).

However, the interpretation of the PCA brings a first trend. Indeed, the samples are clearly separated along the first (horizontal) axis according to the temperature: samples at 22°C are all located on the left (negative coordinates on PC1), whereas samples at 15°C are on the right. This indicates that the effect of temperature is stronger than that of ecotypes because PC1 capture the most important source of variability in the data. The representation of the variables, *i.e.* the transcripts (Figure 7B), is not of great interest at this step; it mainly highlights the need for selection methods to facilitate the interpretation of the results in terms of gene expression level. However, the interpretation of such a plot jointly with the individual plot enables, for instance, identifying over-expressed genes in samples at 15°C: they are located on the right of the variables plot (in the same area as samples at 15°C in the individual plot).

### 5.3. Supervised analysis and variable selection

To illustrate a supervised analysis, we deal with the same data set as before (cell wall transcriptomics for the quantitative block) to discriminate the samples according to the temperature (qualitative block) by performing a PLS-DA analysis. A similar analysis could be made with the ecotype, but interpretation would be more complicated with 5 categories instead of 2 for temperature. Moreover, we have already seen that the temperature effect is the strongest for this data set (Figure 7A). Furthermore, to address the problem of interpretability of the results, we also consider the sparse version of PLS-DA to select the most discriminant genes for the temperature effect. The number of variables to select has to be determined by the user. It depends on the way the potential candidates will be validated. For instance, if validation has to be done through new biological experiments, the number of selected variables must not be too large (about 10). But, if the validation consists in querying a biological database, this number can be higher (about hundreds).

Figure 7 also displays the results of PLS-DA (C, D) and S-PLS-DA (E, F). Individuals plots (Figure 7C, E) and variables plots (Figure7D, F) are interpreted in the same way as PCAs. Individuals plots only use two colors corresponding to the two temperatures. For both PLS-DA and S-PLS-DA, the discrimination between the samples is clear-cut (Figure 7C, E). This result confirms the overriding effect of the temperature. In other words, the variability due to the five ecotypes does not impede from detecting the temperature effect. The result of S-PLS-DA indicates that the discrimination can be observed with only a few genes. Indeed, the difference between PLS-DA and S-PLS-DA relies on the number of genes involved in the discrimination process. The list of the most relevant genes displayed in Figure 7F has to be investigated through for instance functional analysis, but these developments are outside the scope of this article.

These examples of sparse methods highlight the specificity of a supervised analysis: it enables studying the impact of the factors of the experimental design (here the temperature) on the quantitative variables. Thus, the biologist can play with these factors to answer its main biological question and to identify potential future prospects.

### 5.4. Multi-block analyses

Multi-block analyses can address the main purpose of an integrative study by analysing together all the blocks acquired for each sample. As an illustration, we expose the results of a five-block supervised analysis focused on the rosettes, considering phenotypic, cell wall transcriptomics, proteomics and metabolomics as quantitative variables and temperature as the qualitative (or categorical) block.

The statistical relationships between blocks must be defined by the user through a design matrix. This matrix is a square of size [(number of blocks) × (number of blocks)], it is symmetrical and contains values between 0 and 1. A value close to or equal to 1 (respectively 0) indicates a strong relationship (respectively weak or no relationship) between the blocks to be integrated. Fixing the values in the design matrix is crucial and complex because it requires expressing biological relationships as numerical values (*e.g.* can we consider that the link between proteomics and transcriptomics data is stronger than the link between proteomics and metabolomics data?). For the sake of simplicity, 0 and 1 values can be used in a binary point of view: blocks are linked or not. In a supervised context, the values also enable balancing the optimisation between, on the one hand the relationships between quantitative blocks and, on the other hand, the discrimination of the outcome. In our example, we considered a design matrix composed of 0 between blocks to favor the discrimination task rather than the relationships between the blocks. A full design matrix (composed of 1) highlights more clearly relationships between blocks, but can lead to misclassified samples.

The interpretation of a multi-block supervised analysis requires several graphical outputs. Some of them are presented in Figure 8. Figure 8A allows to check whether the correlation between the first components from each data set has been maximized as specified in the design matrix (Tenenhaus et al., 2014). Globally, correlation values are close to 1 and mainly due to the separation of the two categories (22 vs 15°C; because of our design, this matrix favors discrimination). With a full designed matrix, we get higher correlation values but with less separated groups. Regarding the individual plots (Figure 8B), it appears that the discrimination is better for the transcriptomics and proteomics blocks than for the others. The sample plot (Figure 8B) has also to be interpreted regarding the variable plot (Figure 8C). To make the interpretation easier, we present here the results of the sparse version of the multi-block analysis. Therefore, we can identify variables from each block mainly involved in the discrimination according to the temperature. For instance, variables located on the right on the correlation circle plot (Figure 8C) contribute to the discrimination between the samples growing at 22°C because they are also located on the right in the individuals plots (Figure 8B). Another way to display the results is presented in Figure 8D. The clustered image map highlights the profiles of selected variables among the samples. It also includes the results of hierarchical clustering performed jointly on variables and samples. Regarding the samples, the two groups based on temperature are visualized through the dendrogram on the left. However, let us note that the cluster gathering the samples at 15°C can be split into two sub-clusters with the Col ecotype isolated. Regarding the variables, it mainly points out global trends of the behavior of selected variables. The interpretation can then lead to retro analyses to validate potential candidates. This can be done through new statistical analyses as well as new biological experiments (Chawla et al., 2011).

**Figure 8.**
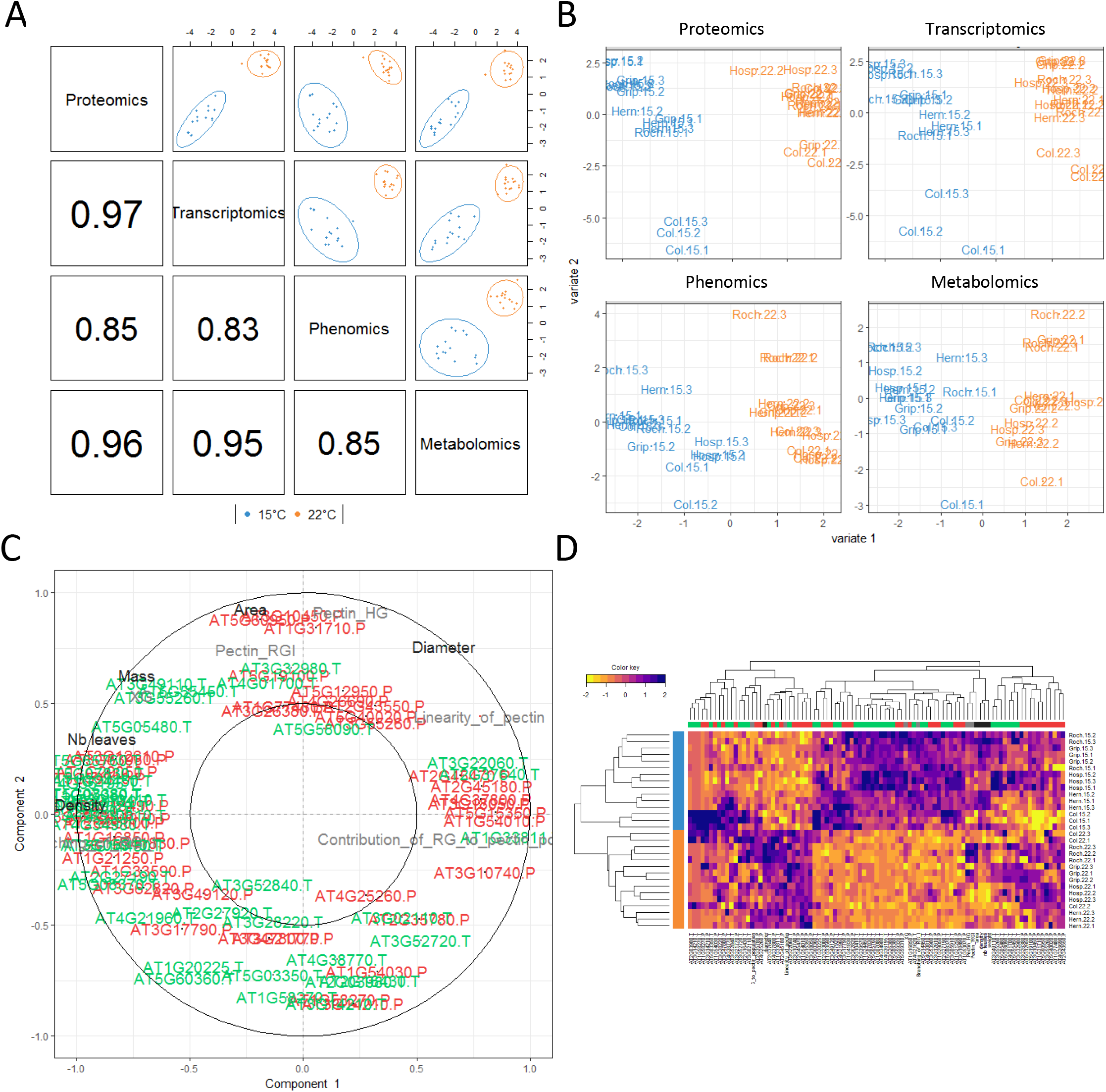
A graphical representation of a multi-block analysis realised on the rosettes of ecotypes grown at 22°C (orange) and 15°C (blue). A) plotDIABLO shows the correlation between components from each data set maximized as specified in the design matrix. B) Individuals plot projects each sample into the space spanned by the components of each block associated to the C) Variables plot that highlights the contribution of each selected variable to each component, D) Clustered image map of the variables (Protein: red; Transcripts: green; Metabolites: grey; Phenotypes: black) to represent the multi-omics profiles for each sample (15°C: blue, 22°C: orange). These plots were obtained using the functions block.splsda(), plotIndiv(), plotVar() and cim() from the mixOmics package (Rohart et al., 2017).

### 5.5. Relevance networks

Another way to interpret the results of a multi-block approach consists in producing relevance networks between variables. On Figure 9A, each selected variable is a node located on a circle. Variables are sorted first according to their block, then depending on their importance in discrimination. An edge links two nodes if their correlation is higher than a threshold subjectively set by the user (we chose 0.9 in Figure 9A).

**Figure 9.**
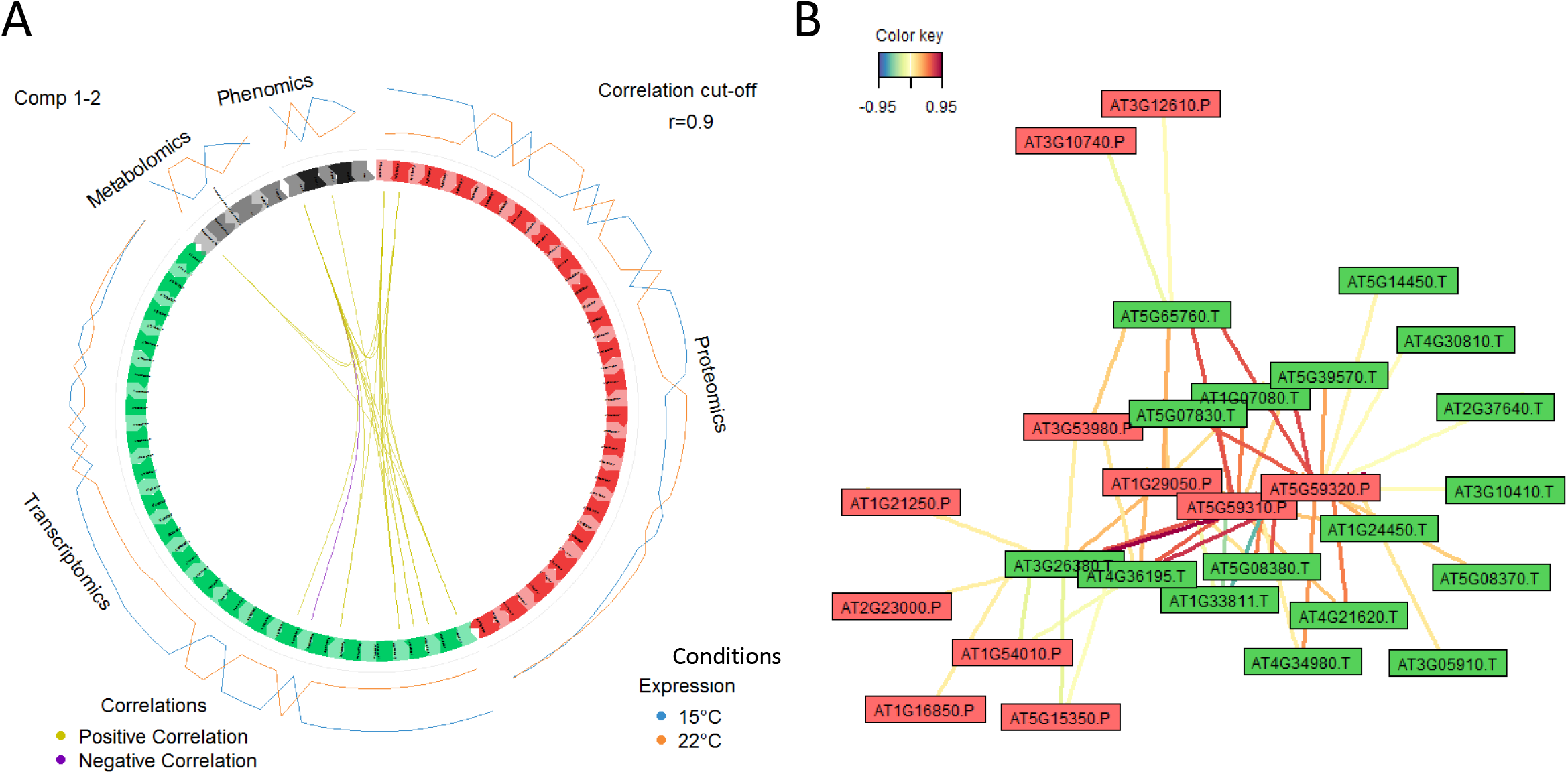
Example of network representation. A) A Circos plot represents the correlations between variables within and between each block (edges inside the circle) and shows the average value of each variable in each condition (line profile outside the circle). B) A network displaying the correlation between the transcriptomics (.T, green) and the proteomics data (.P, red) colored from blue to red according to the color key. These plots were obtained using the functions circosPlot() and network() from the package mixOmics (Rohart et al., 2017).

The correlations are mainly positive and concern a few variables from each block. To complete the interpretation, we focus on another network generated with only two blocks (Figure 9B, cell wall transcriptomics and proteomics). It accentuates the relationships between pairs of proteins and transcripts. The selection of variables is a precious information for the biologist to focus on some of them for validation and draw conclusions in biological terms.

Relevance networks can also be viewed as a first step to modelling as it mimics biological networks and provides clues to address inference networks issues through further dedicated experiments.

## 6. CONCLUSION

In an integrative biology context, the huge quantity of data produced, which can also be heterogeneous, requires adapted and specific statistical methods tentatively summarized in Figure 2. Even if the multi-block approaches can be viewed as the best tool to address a given issue, other more basic standard statistical methods (univariate for instance) must not be omitted. A deep understanding of a biological phenomenon requires a sequence of various approaches to analyse the data. Finally, we consider that each method contributes to a better interpretation of the others as we intended to express it with the schematic view of the protocol as intertwined cycles (Figure 4). The statistical analysis of the large omics data sets can be a never-ending story because each step of the framework provides information. The results presented in this case study could not have been obtained using standard statistical approaches. Actually, it is our global integrative strategy that led us to novel biological results.

## Acknowledgements

The authors are thankful to the Paul Sabatier-Toulouse 3 University and to the *Centre National de la Recherche Scientifique* (CNRS) for granting their work. This work was also supported by the French Laboratory of Excelence project “TULIP”(ANR-10-LABX-41; ANR-11-IDEX-0002-02). HD was supported by the Midi-Pyrénées Region and the Federal University of Toulouse. Thanks to Dr Kim-Anh Lê Cao,Pr Philippe Besse and François Bartolo for their support and help with the graphical outputs and interpretation of the multi-block analysis.

Supplementary Files:

Supplementary Table S1. Toy data set containing 12 observations and 3 variables.

