## Supplemental Table 1 for "A powerful framework for an integrative study with heterogeneous omics data: from univariate statistics to multi-block analysis"

Supplementary Table S1. Toy data set containing 12 observations and 3 variables (1 qualitative group, and 2 quantitative variables, Vx and Vy). The p-values indicated in the last two rows are related to two statistical tests (Wilcoxon rank sum test and Student t test) comparing the observations between Control and Treated separately for each variable.

| **Individual** | **Group** | **Variable Vx** | **Variable Vy** |
|---|---|---|---|
| 1 | Control | 2.0 | 2.0 |
| 2 | Control | 4.5 | 3.5 |
| 3 | Control | 6.0 | 4.3 |
| 4 | Control | 8.0 | 5.1 |
| 5 | Control | 10.0 | 6.0 |
| 6 | Control | 11.0 | 6.5 |
| 7 | Treated | 3.0 | 2.5 |
| 8 | Treated | 5.0 | 3.25 |
| 9 | Treated | 5.5 | 3.3 |
| 10 | Treated | 7.0 | 4.2 |
| 11 | Treated | 8.5 | 4.8 |
| 12 | Treated | 9.0 | 5.0 |
| Test difference Control *vs* Treated | Wilcoxon rank sum test (p-value) | **0.82** | **0.31** |
| | **0.74** | **0.39** | |
